# The fate of melanocytes and the disorganization of basement membrane in a guinea pig model of Rhododendrol-induced chemical vitiligo

**DOI:** 10.1101/2025.09.24.678429

**Authors:** Yasutaka Kuroda, Fei Yang, Lingli Yang, Sylvia Lai, Takuo Yuki, Tetsuya Sayo, Yoshito Takahashi, Daisuke Tsuruta, Ichiro Katayama

**Affiliations:** Department of Pigmentation Research and Therapeutics, Graduate School of Medicine, Osaka Metropolitan University, Osaka, Japan; Cosmetic Products Research, Research and Development, Kao Corporation, Odawara, Japan; Department of Dermatology, Graduate School of Medicine, Osaka Metropolitan University, Osaka, Japan

**Keywords:** rhododendrol, chemical vitiligo, melanocytes, basement membrane, MMP2, Guinea pig model

## Abstract

**Background:** Rhododendrol (RD) is a phenolic compound that was first developed as a skin-lightening agent that occasionally induces skin depigmentation. Although it has been shown that RD induces melanocyte death in vitro, it is still not fully understood why melanocytes are gone by RD in vivo.

**Objective:** This research aimed to investigate how melanocytes are eliminated in the animal model by RD.

**Methods:** On the backs of black guinea pigs (JY-4) with epidermal melanocytes in the basal layer, 30% RD was administered topically three times per day, five days per week. Skin tissues were collected sequentially and histologically analyzed.

**Results:** On day 21, L* values in the RD-applied skin were significantly higher than in the vehicle-applied skin. From day 1 to day 7, the number of TRP1-positive melanocytes and melanin in the basal layer decreased, but no TUNEL-positive melanocytes were identified. On the other hand, an accumulation of melanin was newly found in the dermis. Immunohistochemical staining identified several melanocytes in the upper dermis or spinous layer, away from the basement membrane. An investigation of the epidermal-dermal interface showed a structural anomaly in a portion of the basement membrane with elevated MMP2 expression and increased dermal fibroblasts. The application of the MMP2 inhibitor Ilomastat abolished the basement membrane abnormality by RD.

**Conclusion:** These findings suggest that RD-induced alterations in basement membrane structure may contribute to melanocyte detachment and loss, which is the cause of skin depigmentation not only in RD-induced vitiligo but also in vitiligo.

## Introduction

Chemical-induced vitiligo, also known as contact/occupational vitiligo and chemical leukoderma, is a pigmentary disorder that results in a reduction or loss of skin pigmentation due to repeated exposure to certain chemicals[1, 2]. This condition significantly impacts the quality of life of patients by causing symptoms similar to those of vitiligo. Chemical-induced vitiligo can be induced by many kinds of chemicals, but it is especially reported to be induced by substances with phenolic structure such as monobenzyl ether of hydroquinone (MBEH) and 4-tertiary butyl phenol (4-TBP).

In 2013, the use of cosmetics containing the skin whitening/lightning agent Rhododendrol (RD) (chemical name; 4-[4-hydroxyphenyl]-2-butanol) caused skin depigmentation, with a total of nearly 20,000 cases reported in Japan[3]. Notably, 16% of patients did not demonstrate improvement even 18 months after discontinuing cosmetic use. In vitro studies have demonstrated that RD which possesses a phenolic structure is metabolized by tyrosinase to yield RD metabolites, including RD-quinone and RD-melanin, which induce apoptosis of melanocytes[4-7]. Furthermore, melanocytes exhibiting impaired autophagy and diminished glutathione pools, which possess antioxidant properties, have been observed to be more vulnerable to the toxicity of RD metabolites[6, 8].

Conversely, the use of animal models is imperative for elucidating the percutaneous toxicity of RD on melanocytes. Guinea pigs, mice, and zebrafish have been documented as in vivo models of RD-induced vitiligo[9-12]. We have reported that RD applied continuously to the skin of black guinea pigs or brown guinea pigs pre-irradiated with UV light induced depigmented patches that were clearly brighter than the naive skin in 17 to 21 days[9]. However, in this model, RD application only reduced epidermal melanin and Dopa- or S-100-positive melanocytes. The mechanism by which melanocytes were reduced remains unclear. The present study aims to elucidate the mechanisms by which RD induces apoptosis in melanocytes in vivo and how melanocytes in the basal layer are removed.

## Materials and Methods

### Chemicals

Rhododendrol (RD, 4-[4-hydroxyphenyl]-2-butanol) was kindly provided by Kanebo Cosmetics Inc. (Tokyo, Japan). Ethanol (EtOH) was purchased from Fujifilm Wako Pure Chemical (Osaka, Japan). RD solutions (30w/w%) used in this study were prepared in 50% EtOH (ethanol:water = 1:1). Ilomastat (MedChemExpress, NJ, USA) was dissolved in 50% EtOH.

### Animals

Female black guinea pigs (Kwl:JY-4, 4-6 months old) were purchased from Tokyo Laboratory Animal Science Co., Ltd. (Tokyo, Japan). All animals were kept with ad libitum access to standard feed and water. This study was approved by the Animal Experiment Committee of Osaka Metropolitan University (approval number: 18042).

### Colorimetric measurements

A tristimulus colorimeter (Chromameter, CR-300, Minolta, Tokyo, Japan) was used to evaluate brightness changes of the skin. Color is expressed using the L*a*b* system[13]. In this study, the L* value (brightness) was used, and changes in this parameter are used as an indicator of skin depigmentation[14]. The L* value was measured in each application area (averaging 3 times per point). The mean value of the application area in each animal was obtained from more than 3 animals.

### Histological analysis

For paraffin-embedded sections, the collected skin tissues were fixed with 10% formaldehyde (Fujifilm Wako Pure Chemical) and embedded in paraffin. For Hematoxylin and Eosin (H&E), Fontana-Masson (FM) and immunohistochemical staining, 5-μm thin sections were prepared. H&E staining was performed by deparaffinizing and staining using the Hematoxylin and Eosin solution (Muto pure chemicals, Tokyo, Japan). FM staining was performed by deparaffinizing and staining for melanin using the Fontana-Masson Stain Kit (ab150669, Abcam, Cambridge, UK). Paraffin sections were deparaffinized and heated in Target Retrieval Solution (pH 6) at 95°C for 16 minutes for immunohistochemical staining. Subsequently, the sections were reacted with the primary antibodies (anti-TRP1, HPA000937, 1:200; Sigma-Aldrich or anti-Ki67, M7240, 1:100; Dako) at 4°C overnight. Secondary staining was performed with Alexa Fluor 488 or 555-conjugated anti-rabbit or mouse secondary antibody (1:500; Therma fisher Scientific). Sections were counterstained with Hoechst 33342 (1:500; Therma fisher Scientific). TUNEL staining was performed using the In Situ Cell Death Detection Kit-Fluorescein (1684795, Sigma) according to the manufacturer’s instructions. Positive controls were treated with DNase I for 10 minutes at room temperature.

For frozen sections, the collected skin tissues were fixed with 4% paraformaldehyde (Fujifilm Wako Pure Chemical), washed with PBS(-), dehydrated with 30% sucrose, and embedded in O.C.T. compound (Sakura Finetek, Tokyo, Japan). Subsequently, the sections were reacted with primary antibodies (1:100 dilution) at 4°C overnight. Secondary staining was performed with Alexa Fluor 488 or 555-conjugated anti-rabbit or mouse secondary antibody (1:500; Therma fisher Scientific). The primary antibodies used for frozen sections were as follows: Collagen IV (ab6586, Abcam); Hsp47 (SPA-470, Enzo Life Sciences, NY, USA); MMP2 (ab86607, Abcam). Stained sections were observed under a fluorescence microscope (BZ-X800; Keyence). The quantification of collagen IV staining signal in basement membrane was performed using ImageJ software (ver. 1.52f). The “collagen IV staining intensity per length” was calculated by dividing the “staining intensity of epidermal basement membrane” by the “basement membrane length” at the corresponding site. The relative values were then calculated with the unstimulated condition (Non) in each animal set as 1.

### Electron microscopy analyses

The collected skin tissues were soaked in 2% glutaraldehyde and fixed for 2 hours at room temperature. After washing, they were further fixed in 1% OsO4 for 1 hour at room temperature. After washing, the tissue was dehydrated, embedded in epoxy resin (TAAB EPON, 342-2, Nisshin EM, Tokyo, Japan), thinly sliced, stained with uranyl acetate and lead citrate, and observed with the transmission electron microscope FEI Talos F200C G2 (ThermoFisher Scientific).

### Statistical analysis

The unpaired Student’s t-test was used to evaluate differences in two groups. The Kruskal–Wallis test was used to evaluate differences in diversity among three or more groups, and post hoc analysis was performed with the Dunn test. GraphPad Prism 9 (GraphPad Software Inc., San Diego, CA, USA) was used for statistical analysis. A p-value <0.05 was considered statistically significant.

## Results

### Induction of RD-induced depigmentation in black guinea pigs

To analyze sequential changes in the epidermal melanocytes, 30 (w/w)% RD or vehicle (50 [v/v]% EtOH/Water) was topically applied to a shaved lateral abdomen (Fig. 1a). Treatment frequency was three times a day, 5 days a week for 14 weeks (98 days). After repeated application of RD to JY-4 black guinea pigs, clear depigmentation spots were observed around day 14 (Fig. 1b). The L* value, an index of brightness, was significantly higher on day 21 and reached a plateau on day 28 (Fig. 1c). Next, we analyzed the histological distribution of melanin and melanocytes in the skin during the depigmentation process (Fig. 2). The results showed that melanin was gradually decreased from day 4, mainly in the basal layer, and almost no melanin could be observed in the epidermis by day 49 (Fig. 2b). For TRP1-positive melanocytes in the epidermis, the number of cells was already markedly decreased on day 1, and many areas were almost entirely absent even on day 4, and almost all areas were not identified on day 49 (Fig. 2c). During this period, no significant mononuclear cell infiltration such as lymphocytes was observed by HE staining (Fig. 2a). Thus, the application of RD resulted in the disappearance of most TRP1-positive melanocytes within a few days without excessive inflammation.

**Figure 1.**
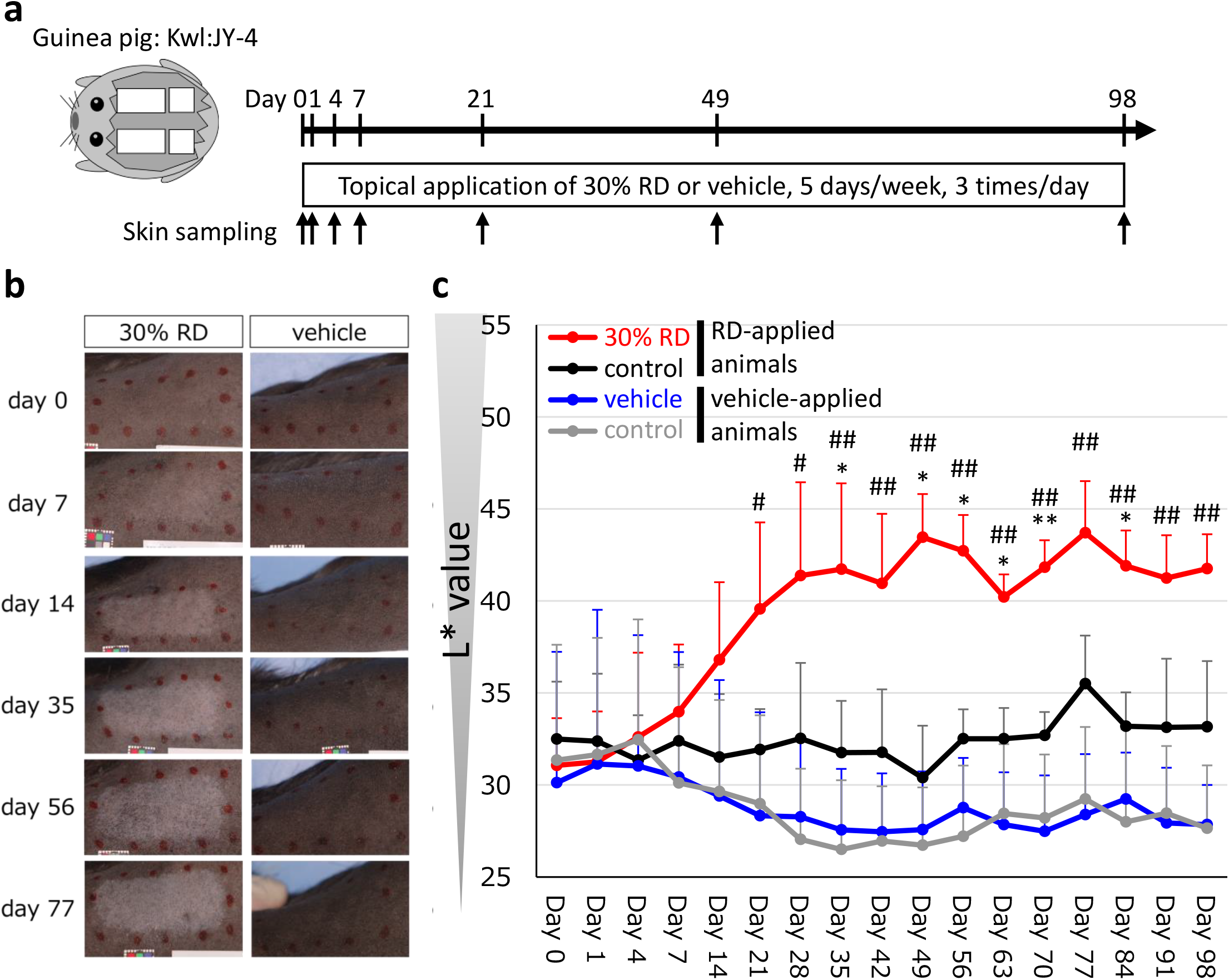
Induction of skin depigmentation by RD in black guinea pigs. (a) Experimental design. (b) The skin color of RD or vehicle treated skin. (c) Time course of L* values of RD or vehicle treated skin or untreated skin (n = 3). Data in are presented as means ± SD. *p <.05, **p <.01 (30%RD vs control in RD-applied animals), #p <.05, ## p <.01 (30%RD vs vehicle); difference between the indicated two groups by unpaired Student’s t-test.

**Figure 2.**
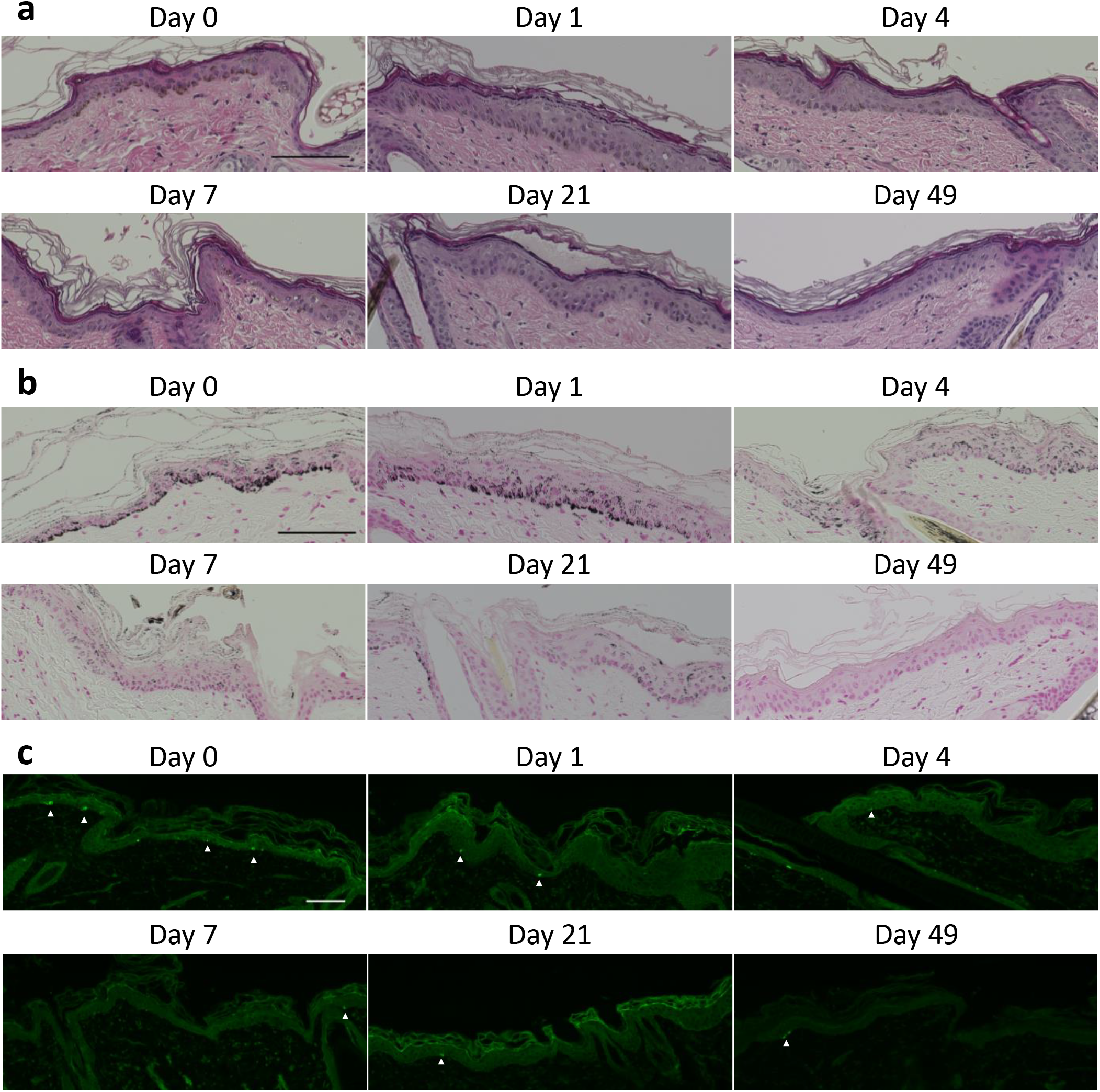
Histological examination and the distribution of melanin and melanocytes in the skin during the depigmentation process. (a) HE staining images, (b) Fontana-Masson staining images and (c) immunohistochemical images of TRP1 are presented. Scale bar, 100 μm.

### Melanocyte loss is caused by rather cell death than detachment from basement membrane

As mentioned above, depigmentation caused by RD was observed around day 14, thus it is hypothesized that alterations of melanocyte behavior should have occurred earlier. Therefore, to reveal the presence of apoptotic melanocytes by day 14, we examined the presence of apoptotic cells in the melanin-containing cells of the basal layer of the epidermis by TUNEL staining. The results showed that TUNEL-positive cells in the basal layer of the epidermis were absent from day 0-14, while positive cells were occasionally observed in the dermis (Fig. 3). There is negligible apoptosis of epidermal melanocytes due to RD application on days 1-14 in this model.

**Figure 3.**
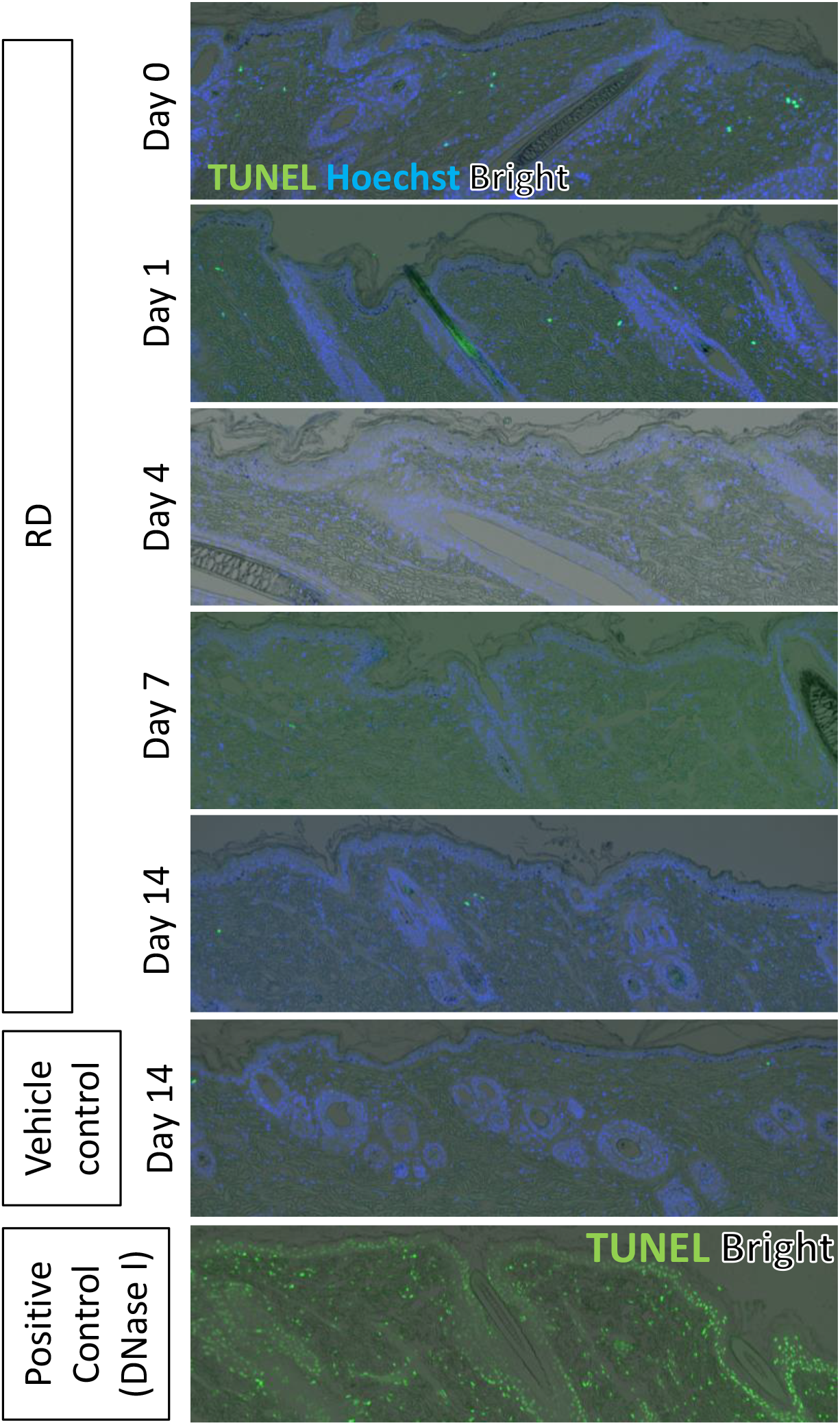
TUNEL staining (in green) images along with bright field and nuclear staining images with Hoechst33342 (in blue) in the skin during the depigmentation process. Scale bar, 100 μm.

Careful observation of histological images of the skin treated with RD for 1-4 days revealed the presence of melanin in the upper dermal layer (Fig. 4a). Melanin in the upper dermis was almost absent on day 0. In addition, immunohistochemistry of TRP1 revealed that melanocytes are usually located in the basal layer, however, in RD-applied skin, melanocytes were also present in the spinous layer of the epidermis or in the upper dermis, away from the basement membrane (Fig. 4b). In this case, the marker of cell proliferation, Ki67, was negative, along with nearby melanocytes. Electron microscopy also revealed detached melanocytes away from the basal layer, melanocytes in the dermis, and melanocyte-like cells that may have died in the dermis (Fig. 4c). Thus, it was suggested that melanocytes could be detached from the basement membrane by RD, floating to the stratum corneum, or falling into the upper dermis. Further observation of the basement membrane by electron microscopy revealed (1) a decrease in electron density of the lamina densa and (2) disappearance of the lamina lucida in part of basement membranes due to RD application (Fig.5a). Furthermore, type IV collagen, a major component of the basement membrane, was partially undetected on day 4 by immunohistochemical staining (Fig. 5b). These findings suggest that RD could triggered structural aberrations within the basement membrane.

**Figure 4.**
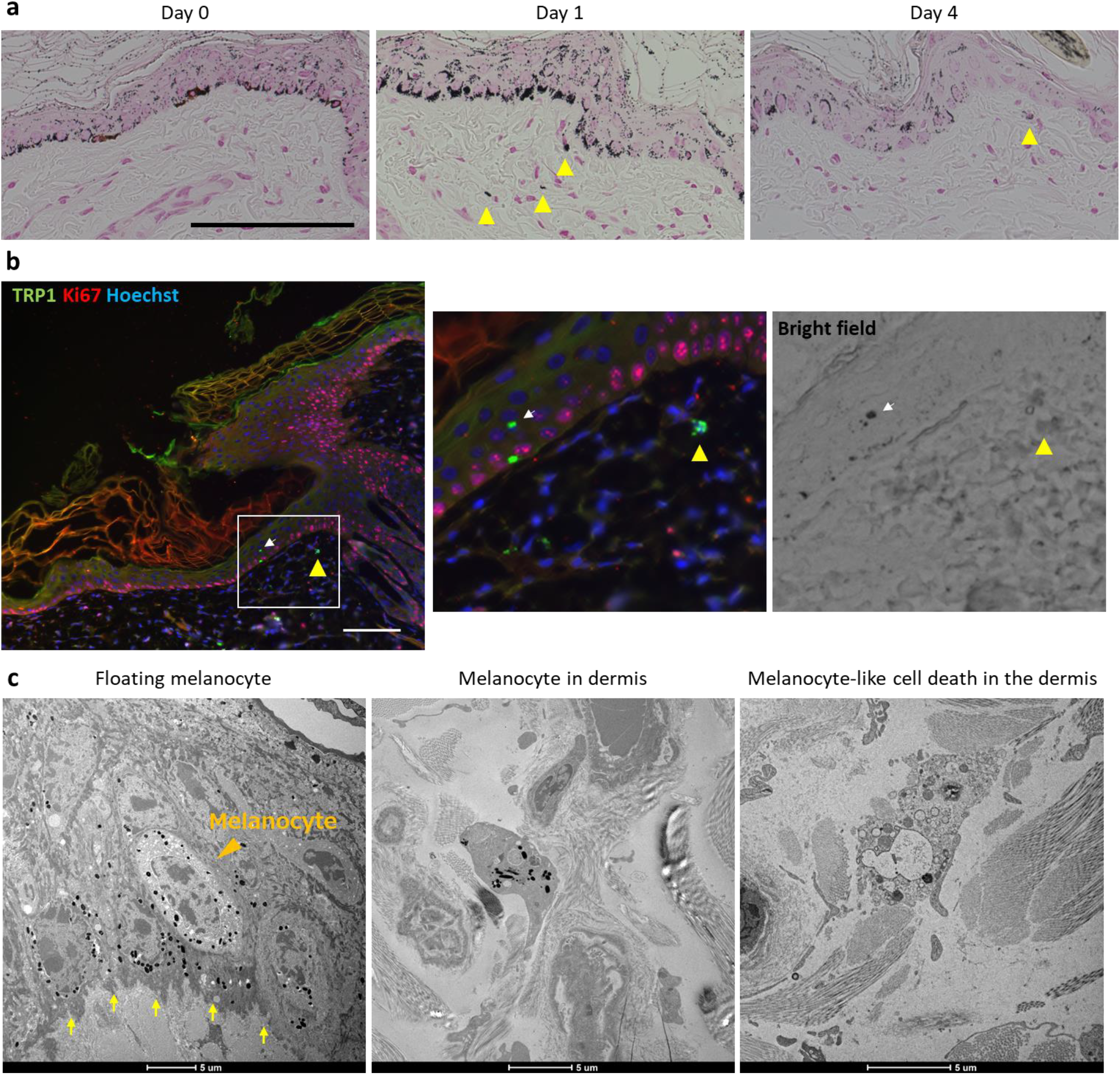
Distribution of melanin and melanocytes in the skin during the early depigmentation process. (a) Fontana-Masson staining images and (b) immunohistochemical images of TRP1 (in green) and Ki67 (in red) along with nuclear staining with Hoechist33342 (in blue) are presented. The area in the white rectangle is shown enlarged in the right panels with the corresponding bright-field image. Scale bar, 100 μm. (c) Representative ultrastructural images of ultrathin sections. Scale bar, 1 μm.

**Figure 5.**
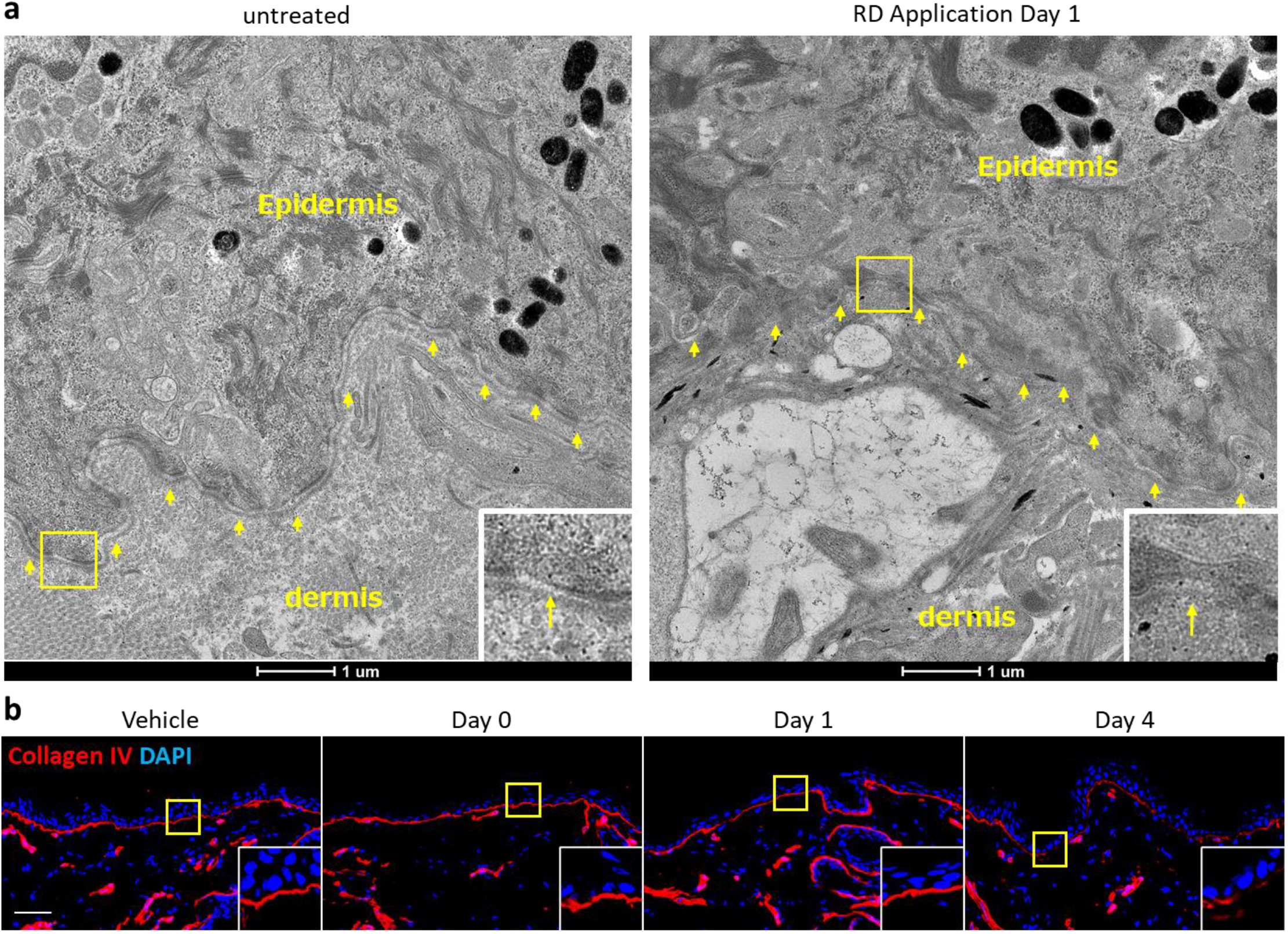
Partial structural aberrations with the basement membrane in RD-applied skin in guinea pigs. (a) Representative ultrastructural images of ultrathin sections. The areas in the yellow rectangles are shown enlarged in the lower right corner of the images. Yellow arrows indicate the basement membrane. Scale bar, 1 μm. (b) Immunohistochemical images of type IV collagen (in red) along with nuclear staining with DAPI (in blue) are presented. The areas in the yellow rectangles are shown enlarged in the lower right corner of the images. Scale bar, 100 μm.

### RD application induced anormal basement membrane accompany with the elevated number of fibroblasts and increased MMP2 expression which associated with type IV collagen expression

Since our previous report showed that MMP2 by dermal fibroblasts induces basement membrane disorganization in vitiligo[15], we investigated the possibility that basement membrane damage by RD also occurs along the fibroblast-MMP2 axis. Frozen sections from the RD-treated skin for 1-4 days were stained with anti-HSP47 (fibroblast marker) antibodies and anti-MMP2 antibodies. HSP47-positive fibroblasts were dramatically increased on days 1 and 4 (Fig. 6a). Similarly, MMP2 expressions were markedly increased on days 1 and 4 in the epidermis and dermis (Fig. 6b). Besides, to investigate the involvement of MMP2 in the basement membrane, Ilomastat, a potent inhibitor of MMP2, was applied for 4 days starting 1 day prior to RD application, and RD was also applied concurrently for 3 days (Fig. 6c). On day 4, the skin was collected and frozen thin section samples were stained with anti-type IV collagen antibody. The RD application led to a substantial decrease in the intensity of type IV collagen staining of the basement membrane (Fig. 6d,e). However, the simultaneous application of Ilomastat effectively counteracted this decrease in staining intensity. Accordingly, these findings suggest that RD induces the accumulation of fibroblasts and MMP2, and that at least MMP2 contributes to the degradation of basement membranes (Fig. 6f).

**Figure 6.**
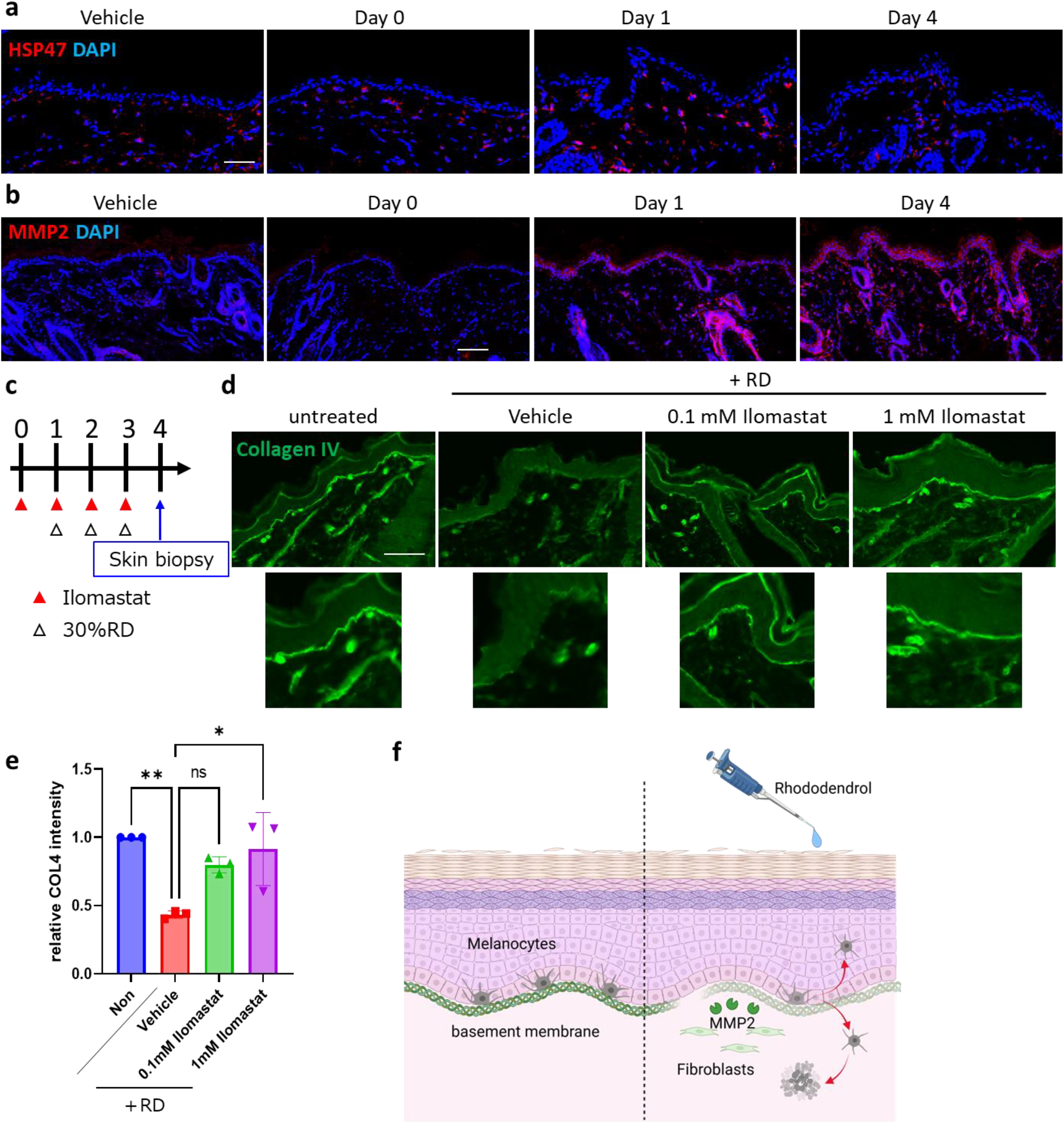
MMP2-dependent anormal basement membrane is induced by RD accompany with the accumulation of fibroblasts and elevated MMP2 expression. (a) Immunohistochemical images of HSP47 (in red) along with nuclear staining with DAPI (in blue) are presented. (b) Immunohistochemical images of MMP2 (in red) along with nuclear staining with DAPI (in blue) are presented. Scale bar, 100 μm. (c) Experimental design. (d) Immunohistochemical images of type IV collagen are presented. the enlarged details are shown in the corresponding lower panels. Scale bar, 100 μm. (e) Relative signals of anti-type IV collagen antibody within basement membrane are quantified. Data in are presented as means ± SD. *p <.05, **p <.01; difference in diversity among three or more groups by the Kruskal–Wallis test, and post hoc analysis was performed with the Dunn test. (f) Summary illustration of the possible response of epidermal melanocytes to RD in guinea pig models.

## Discussion

In this study, we examined the process of melanocyte loss in a guinea pig model of RD-induced vitiligo and proposed that one of the factors contributing to melanocyte loss is not apoptosis in the epidermal basal layer of melanocytes, but rather detachment of melanocytes from the basement membrane, which serves as scaffold for melanocytes. Furthermore, structural abnormalities in the basement membrane by MMP2 were observed during the process of melanocyte detachment from the basement membrane. These observations suggest a pattern of melanocyte loss in which the basement membrane is disrupted by RD, leading to the detachment of melanocytes from the basement membrane and subsequent disappearance.

In reported animal models of RD-induced vitiligo, the mechanism of melanocyte loss has been investigated only in the K14-Kitl transgenic mice model[16]. In this model, the expression of DDR1 and E-Cadherin in the epidermis, which are involved in melanocyte adhesion, temporally decreased on day 7 after continuous application of RD and detached melanocytes were observed. However, the melanocytes dropped into the dermis have not been reported. Therefore, the guinea pig model may be appropriate for analysis of dropped melanocyte to the dermis. In the guinea pig model, the expression of E-Cadherin was also examined, but no significant change was observed (Fig. S1). The underlying mechanisms of melanocyte loss by RD may exhibit common and distinct pathways in both the mouse and guinea pig models due to differences in species-specific altered proteins and skin tissue structure.

The major structural components of basement membrane are type IV collagen, laminin, nidogen, and the heparan sulfate proteoglycans perlecan and agrin[17]. This study revealed the presence of structural abnormalities in the basement membrane of RD-applied skin. In previous studies, type IV collagen staining in patients with vitiligo was different from that in healthy subjects, with the abnormal basement membrane morphologies, including thickening, severe disruption, and multilayering[15]. Furthermore, the number of HSP47-positive fibroblasts increased in the dermis, and most fibroblasts were MMP2 positive, suggesting that fibroblast-derived MMP2 is degrading the basement membrane and atypical remodeling. In this guinea pig model of RD-induced vitiligo, partial abnormalities of type IV collagen in the basement membrane, increased numbers of dermal fibroblasts, and increased expression of MMP2 were observed. The present study demonstrates that RD-induced basement membrane abnormality (decreased type IV collagen expression) is reversed by concomitant use of Ilomastat, a potent MMP2 inhibitor. MMP2 has a wide range of degrading activities, including type IV, V, VII, X and XI collagens, elastin, fibronectin and laminins[18]. Thus, although MMP2 may affect laminins in addition to the type IV collagen, it was suggested that MMP2 plays a role in RD-induced vitiligo pathogenesis and that its inhibition may be a critical component of chemical-induced vitiligo treatment. On the other hand, it has been reported that RD is converted to a toxic quinone form by tyrosinase in melanocytes, which cause melanocyte toxicity[4], but the reason why MMP2 expression is elevated in fibroblasts that lack tyrosinase is still unclear and requires further investigation.

There were no reports on the phenomenon of melanocyte detachment in the patients with RD-induced vitiligo. It is conceivable that the melanocyte-detachment phenomenon could not be substantiated because the specimens of the patients that had been examined were specimens in which the induction of depigmentation had been achieved. Although melanocytes themselves have not been reported to fall into the dermis, melanophages, which are melanin-phagocytized cells, have been reported to be observed at high frequency in the lesion and peri-lesion of the patients with RD-induced vitiligo. Therefore, it is plausible that detached melanocytes may have died in the dermis and been phagocytosed by macrophages.

This study proposes a novel pathway through which melanocytes are eliminated by RD. The MMP2-basement membrane abnormality pathway has been postulated to play a pivotal role in vitiligo, and the RD-induced vitiligo model may facilitate a more profound comprehension of not only RD-induced vitiligo but also non-segmental vitiligo. A critical consideration for future research is the observation of abnormalities in the basement membrane structure in patients with chronic RD-induced vitiligo.

## Abbreviations

RD: Rhododendrol
EtOH: Ethanol
TRP1: Tyrosinase related protein 1
TUNEL: terminal deoxynucleotide transferase dUTP nick end-labeling
MMP2: Matrix metalloproteinase 2
HE: Hematoxylin and Eosin
FM: Fontana-Masson
HSP47: Heat shock protein 47

## Acknowledgments

We appreciate the cooperation and help from our secretary, Kumiko Mitsuyama. Electron microscopy analyses were performed in Research Support Platform of Osaka Metropolitan University Graduate School of Medicine

## Supporting information

**Supplemental Figure 1** Immunohistochemical images of E-Cadherin and TRP1 in the skin during the early depigmentation process.

## Highlights

- Rhododendrol caused melanocytes to disappear from the epidermal basal layer of black guinea pigs in a few days.
- Melanocyte removal by Rhododendrol was not apoptosis of melanocytes but rather detachment of melanocytes from the basement membrane.
- Rhododendrol increased fibroblasts in the dermis, induced MMP2 expression, and aberrant basement membrane structure in an MMP2-dependent manner.

## Notes

**Conflicts of interest:** The authors have no conflict of interest to declare.

**Funding:** This work was supported by the KAO Corporation.

### Competing Interest Statement

The authors have declared no competing interest.

## References

[1] J.E. Harris, Chemical-Induced Vitiligo, Dermatol Clin 35(2) (2017) 151–161.

[2] R.E. Boissy, P. Manga, On the Etiology of Contact/Occupational Vitiligo, Pigment Cell Research 17(3) (2004) 208–214.

[3] K. Matsunaga, K. Suzuki, A. Ito, A. Tanemura, Y. Abe, T. Suzuki, et al., Rhododendrol-induced leukoderma update I: Clinical findings and treatment, The Journal of Dermatology 48(7) (2021) 961–968.

[4] S. Ito, K. Wakamatsu, Biochemical Mechanism of Rhododendrol-Induced Leukoderma, Int J Mol Sci 19(2) (2018) 552.

[5] M. Sasaki, M. Kondo, K. Sato, M. Umeda, K. Kawabata, Y. Takahashi, et al., Rhododendrol, a depigmentation-inducing phenolic compound, exerts melanocyte cytotoxicity via a tyrosinase-dependent mechanism, Pigment Cell & Melanoma Research 27(5) (2014) 754–763.

[6] L. Yang, F. Yang, M. Wataya-Kaneda, A. Tanemura, D. Tsuruta, I. Katayama, 4-(4-Hydroroxyphenyl)-2-butanol (rhododendrol) activates the autophagy-lysosome pathway in melanocytes: Insights into the mechanisms of rhododendrol-induced leukoderma, Journal of Dermatological Science 77(3) (2015) 182–185.

[7] S. Inoue, I. Katayama, T. Suzuki, A. Tanemura, S. Ito, Y. Abe, et al., Rhododendrol-induced leukoderma update II: Pathophysiology, mechanisms, risk evaluation, and possible mechanism-based treatments in comparison with vitiligo, The Journal of Dermatology 48(7) (2021) 969–978.

[8] M. Kondo, K. Kawabata, K. Sato, S. Yamaguchi, A. Hachiya, Y. Takahashi, et al., Glutathione maintenance is crucial for survival of melanocytes after exposure to rhododendrol, Pigment Cell & Melanoma Research 29(5) (2016) 541–549.

[9] Y. Kuroda, Y. Takahashi, H. Sakaguchi, K. Matsunaga, T. Suzuki, Depigmentation of the skin induced by 4-(4-hydroxyphenyl)-2-butanol is spontaneously re-pigmented in brown and black guinea pigs, The Journal of Toxicological Sciences 39(4) (2014) 615–623.

[10] Y. Abe, K. Okamura, M. Kawaguchi, Y. Hozumi, H. Aoki, T. Kunisada, et al., Rhododenol-induced leukoderma in a mouse model mimicking Japanese skin, Journal of Dermatological Science 81(1) (2016) 35–43.

[11] M. Iida, A. Tazaki, Y. Deng, W. Chen, I. Yajima, L. Kondo-Ida, et al., A unique system that can sensitively assess the risk of chemical leukoderma by using murine tail skin, Chemosphere 235 (2019) 713–718.

[12] M. Hayazaki, O. Hatano, S. Shimabayashi, T. Akiyama, H. Takemori, A. Hamamoto, Zebrafish as a new model for rhododendrol-induced leukoderma, Pigment Cell & Melanoma Research 34(6) (2021) 1029–1038.

[13] A.R. Robertson, The CIE 1976 color - difference formulae, Color Research & Application 2(1) (1977) 7–11.

[14] J.C. Seitz, C.G. Whitmore, Measurement of erythema and tanning responses in human skin using a tri-stimulus colorimeter, Dermatology 177(2) (1988) 70–75.

[15] F. Yang, L. Yang, Y. Kuroda, S. Lai, Y. Takahashi, T. Sayo, et al., Disorganisation of basement membrane zone architecture impairs melanocyte residence in vitiligo, The Journal of Pathology 264(1) (2024) 30–41.

[16] Y. Abe, Y. Hozumi, K. Okamura, T. Suzuki, Expression of discoidin domain receptor 1 and E-cadherin in epidermis affects melanocyte behavior in rhododendrol-induced leukoderma mouse model, The Journal of Dermatology 47(11) (2020) 1330–1334.

[17] A. Pozzi, P.D. Yurchenco, R.V. Iozzo, The nature and biology of basement membranes, Matrix Biology 57-58 (2017) 1–11.

[18] T. Shiomi, V. Lemaître, J. D’Armiento, Y. Okada, Matrix metalloproteinases, a disintegrin and metalloproteinases, and a disintegrin and metalloproteinases with thrombospondin motifs in non-neoplastic diseases, Pathology International 60(7) (2010) 477–496.

