## Supplementary figure 1 for "The fate of melanocytes and the disorganization of basement membrane in a guinea pig model of Rhododendrol-induced chemical vitiligo"

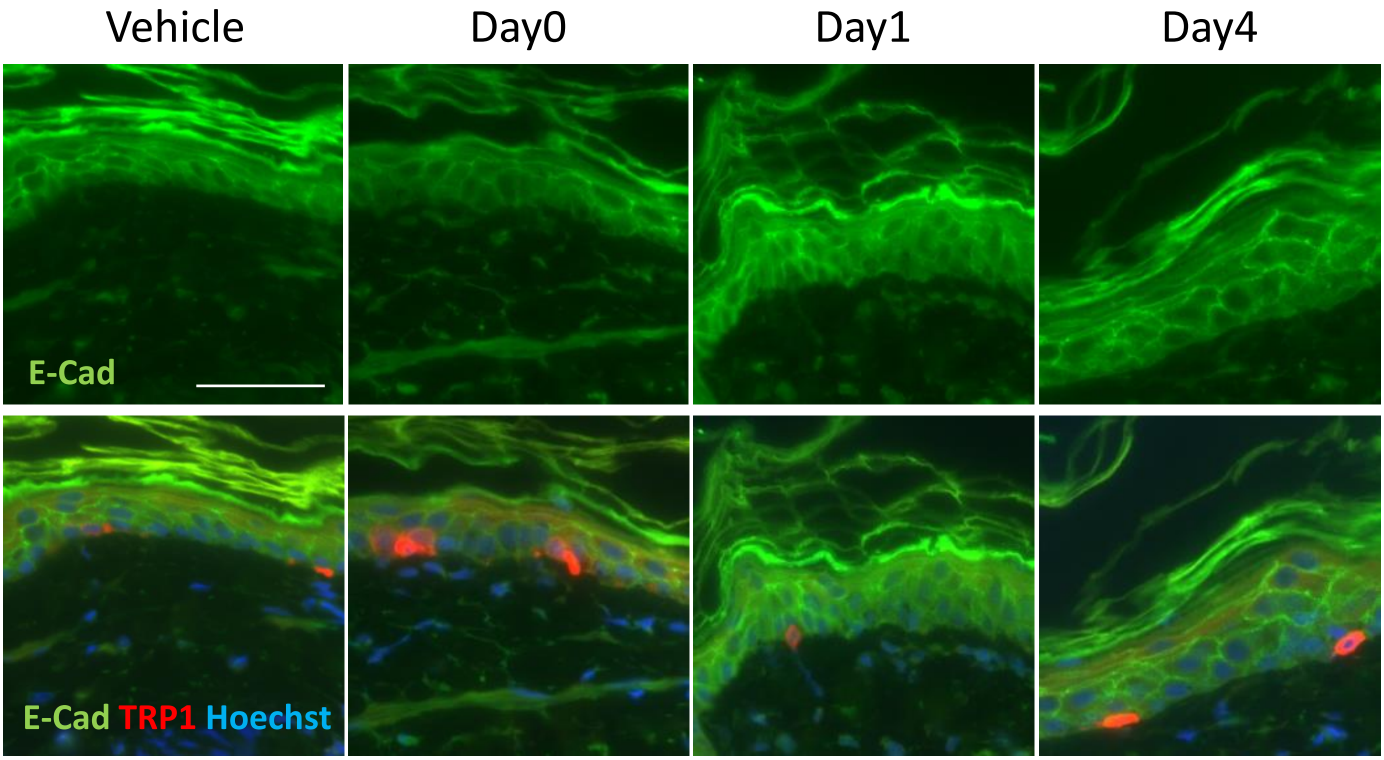


### Supplemental Fig. 1

Immunohistochemical images of E-Cadherin and TRP1 in the skin during the early depigmentation process. Scale bar: 50 μm.
